# Possible exposure to unidentified coronaviruses in roe deer (*Capreolus capreolus*) populations suggested by SARS-CoV-2 serological investigation in France

**DOI:** 10.1101/2025.02.19.639030

**Authors:** Grégoire Perez, Lucas D. Lalande, Vincent Legros, Angéli Kodjo, Hélène Verheyden, Vincent Bourret, Nicolas Cèbe, Yannick Chaval, Paul Revelli, Maryline Pellerin, Jean-François Lemaître, Carole Peroz, Benjamin Rey, François Débias, Rebecca Garcia, Gilles Bourgoin, Emmanuelle Gilot-Fromont

**Affiliations:** Université de Lyon, Université Lyon 1, CNRS, Laboratoire de Biométrie et Biologie Evolutive UMR 5558, F-69622 Villeurbanne, France; CIRAD, UMR ASTRE, Conakry, Guinée; ASTRE, CIRAD, INRAE, Université de Montpellier, 34398 Montpellier, France; CERFIG, Université Gamal Abdel Nasser de Conakry, Conakry, Guinée; Université de Toulouse, Institut National Universitaire Jean-Francois Champollion, BTSB-UR 7417, Place de Verdun, 81000 Albi, France; Université de Lyon, Université Lyon 1, CNRS, Centre International de Recherche en Infectiologie, Inserm U1111 UMR5308, ENS de Lyon, Lyon, France; Université de Lyon, VetAgro Sup, Laboratoire des Leptosires et d’Analyses Vétérinaires, 69280 Marcy L’Etoile, France; Université de Toulouse, INRAE, CEFS, Castanet Tolosan, 31326, France; LTSER ZA PYrénées GARonne, Auzeville-Tolosane, 31320, France; Office Français de la Biodiversité, Direction de la Recherche et de l’Appui Scientifique, Service Conservation et Gestion Durable des Espèces Exploités, 52210 Châteauvillain, France; Université de Lyon, VetAgro Sup, 69280 Marcy l’Etoile, France

**Keywords:** COVID-19, wild ungulates, cervids, spill-over, spill-back, zoonoses

## Abstract

The risk of viral transmissions from domestic/wild animals to humans is a major public health concern. Humans can also transmit viruses back to domestic and wild animals, acting as reservoir for virus maintenance and sources of epidemic diseases re-emergence. The SARS-CoV-2, causing COVID-19, likely originated from wildlife and has been evidenced to transmit from humans to captive, domestic and wild animals. White-tailed deer (*Odocoileus virginianus*) show high SARS-CoV-2 prevalence following human contamination, suggesting they could act as an emerging virus reservoir. We completed recent research on European cervid species by investigating whether SARS-CoV-2 had emerged in longitudinally-monitored European roe deer (*Capreolus capreolus*) populations in direct contact with humans in France. We performed indirect tests (serological ELISAs and seroneutralization) on sera collected before and after the virus emergence in humans, and direct RT-PCR tests on nasal swabs collected in 2022. We also investigated the virus exposure and prevalence in three other cervid species. ELISA tests were positive for 2.20% of sera, pre- and post-pandemic, but seroneutralization and PCR tests were negative. Although one population showed increased seroprevalence post-2020, results suggest that SARS-CoV-2 has not emerged in those populations, and that ELISA cross-reaction with one or several unidentified circulating coronaviruses is possible.

## 1. Introduction

The potential transmission of viral infections inducing an epidemic from livestock or wildlife to humans is of high concern for human health worldwide, as vertebrates constitute the main source of human emerging diseases [1]. Conversely, human pathogens can be transmitted to domestic or wild animal species, potentially inducing new epizooties in these species (*e.g.* human coronavirus OC43 outbreak in wild chimpanzees [2]). Thus, there is a need to identify pathogens features (*e.g.* mutation rate, transmission route) and ecological contexts (*e.g.* species interactions, habitat use) that promote such events.

Coronaviruses are hosted by many mammal species and can be transmitted from one species to another, including humans [2–4]. The COVID-19 pandemic, due to the Severe Acute Respiratory Syndrome Coronavirus 2 (SARS-CoV-2), most likely originated from wildlife, in this case bats [*e.g.* 5,6]. Since the beginning of the COVID-19 pandemic, transmission events have been documented from humans back to domestic animals, for example farmed minks (*Neovison vison*) [7,8], domestic cats (*Felis catus*) and dogs (*Canis lupus familiaris*) [9,10] or wild felids kept in zoos [11]. These first observed transmission events occurred in domestic, captive or intensive farming contexts and could be explained by elevated rates of direct contacts between humans and animals in such specific contexts. However, multiple events of transmission of SARS-CoV-2 from human to free-living white-tailed deer (*Odocoileus virginianus*) have also been documented in North America. These events were followed by transmissions within local white-tailed deer population, resulting in populations displaying high prevalence levels which were up to 40% by seroprevalence, and 82.5% and 70% in retropharyngeal lymph node and nasal swabs tested by RT-PCR, respectively [12–15]. Eventually, three cases of transmission from the newly infected white-tailed deer populations to humans were detected, suggesting that the white-tailed deer may become an emerging virus reservoir in North America [15].

Similar contamination pathways can be expected in Europe, as several species of cervids are highly abundant and can be exposed to viral transmission due to close contact with humans (*e.g.* zoos, parks, research stations). Following this rationale, several recent studies examined whether European cervid species can act as reservoir for the SARS-CoV-2 virus. In most cases, deer populations were found to be uninfected by the virus (UK - chinese water (*Hydropotes inermis*), fallow deer (*Dama dama*), red deer (*Cervus elaphus*), muntjac (*Muntiacus reevesi*), roe deer (*Capreolus capreolus*), sika deer (*Cervus nippon*), red/sika hybrid [16]; Poland – red deer [17]; Germany-Austria – fallow deer, red deer, roe deer [18]; Germany – fallow deer, red deer, roe deer [19]). However, first cases of SARS-CoV-2 seropositivity have been more recently observed in fallow deer in Ireland [20], in free-living fallow and red deer suburban populations in Spain [21], and in a British fallow and red deer park population [22], suggesting that the host tropism of SARS-CoV-2 may change as new variants emerge.

In this context, we extended current European research on cervid species by investigating whether SARS-CoV-2 had emerged in roe deer (*Capreolus capreolus*) populations subject to hunting or direct handling as part of capture-mark-recapture programs in France. Among European cervid species, the roe deer is particularly interesting. The species occupies a large variety of habitats, including agricultural areas with proximity to domestic species and human activities [23], making roe deer potentially exposed to human-borne viruses. The species can also display a gregarious behaviour in winter in open landscapes [24], favouring disease transmission. Furthermore, the hunting regulations and the absence of large predator during the last 50 years resulted in high abundance of roe deer all over Europe. Between 1984 and 2005, the number of roe deer in Europe was estimated to increase from 6.2 to 9.5 million and hunting grew from 1.7 to 2.7 million of hunted individuals yearly [25]. In France, the roe deer is widely hunted (c. 600,000 individuals officially hunted in 2022, i.e. ten times the number hunted 50 years ago) [26]. The rate of potential contact with humans, one of the potential routes of transmission of the virus from humans to cervids, and particularly roe deer, is therefore particularly high in France [27].

To investigate SARS-CoV-2 presence, we benefited from longitudinal studies of four free-living (including three enclosed) and one captive populations of roe deer. Because of annual captures, roe deer from these populations are particularly exposed to human viruses’ transmission. Moreover, the continuous follow-up of the same individuals from 2010 to 2022 was an opportunity to detect virus arrival in populations. The virus was searched for by means of either direct PCR testing (on individuals captured in 2022) or indirect serological testing to assess exposure before and after SARS-CoV-2 emergence in human in France in 2020. We further investigated exposure and presence of the virus in other cervid samples collected in 2022 during capture or hunting operations: red deer, fallow deer and sika deer (*Cervus nippon*).

## 2. Material and methods

### 2.1 Roe deer populations

Roe deer sera were sampled from five populations: Trois-Fontaines (TF), Chizé (CH), Aurignac (AU), Gardouch (GA) and Domain of Praillebard (DP). Among these populations, three are hunted wild populations: TF, CH and AU (Figure 1, red squares). TF is a 1,360 ha fenced forest in north-eastern France (48°43’N, 4°55’E) hosting an enclosed wild roe deer population; CH is a 2,614 ha fenced Integral Biological Reserve (unmanaged forest) since 2006 located in western France (46°05’N, 0°25’W) hosting also a closed wild roe deer population; AU is an open study area covering about 7,500 ha in south-western France (43°16’N, 0°53’E) including forests and agricultural landscapes with free-ranging roe deer.

**Figure 1.**
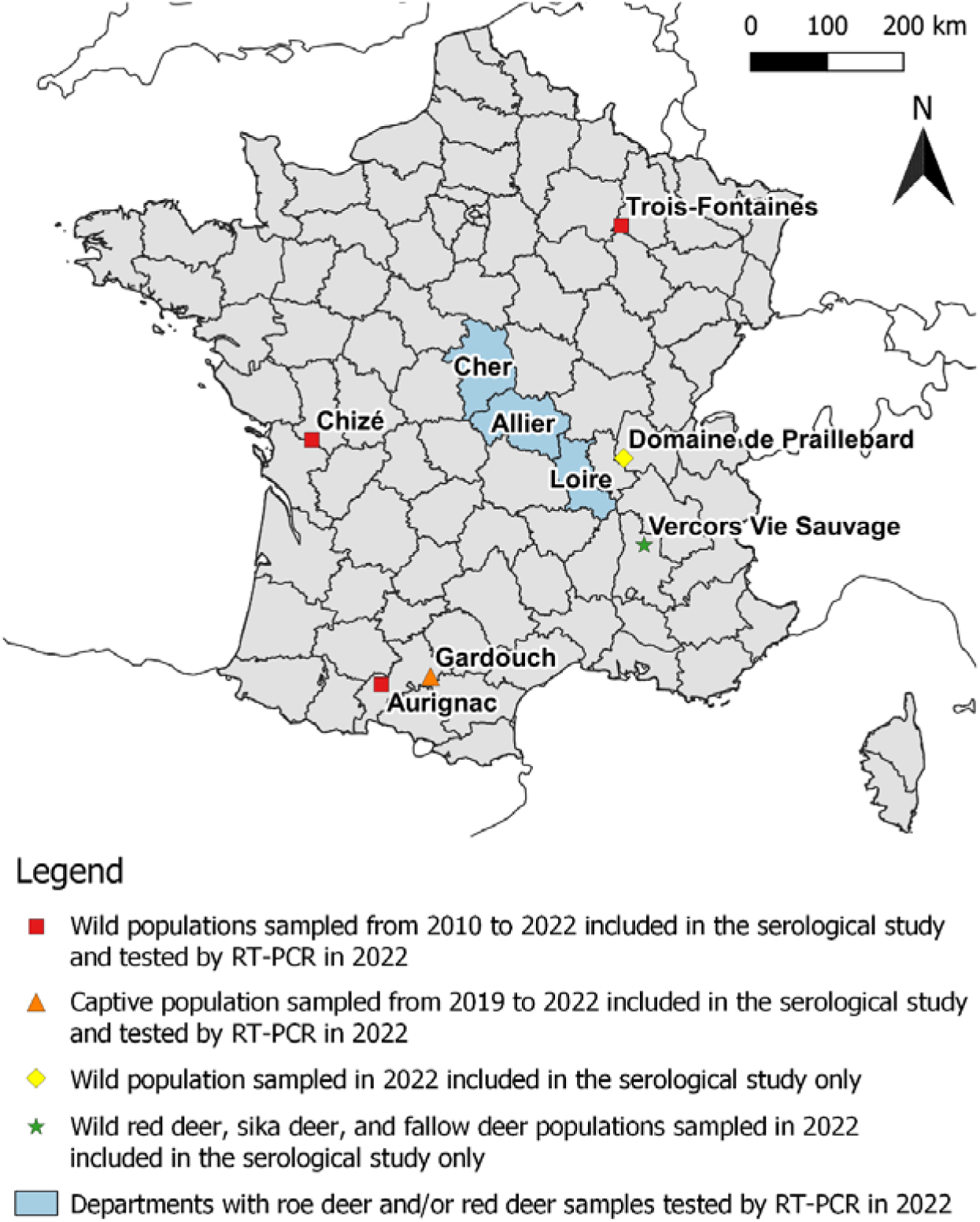
Locations of deer populations sampled for the sera used in the SARS-CoV-2 serological study and the nasal swabs used in the SARS-CoV-2 RNA RT-PCR screening.

Captive roe deer were sampled for their sera in GA (Figure 1, orange triangle), an experimental station in southern France (43°22’, 1°40’E) hosting captive roe deer in small 0.5 ha enclosures or semi-captive roe deer in a large enclosure composed of 9 ha of forest and 5 ha of meadows.

Roe deer sera were also sampled at the DP (Figure 1, yellow diamond), a private fenced area of 148 ha in eastern France (45°57’N, 4°55’E) hosting a small roe deer population introduced from TF and CH in 2019-2020. Sera sampling is summarized in Figure 1 and Table 1.

**Table 1.** Summary of the sampling scheme and the SARS-CoV-2 serological results for roe deer (*Capreolus capreolus*) and other cervids for pre- (*i.e*.. 2010-2020) and post- (i.e. 2021-2022) COVID-19 emergence, set in March 2020. Number of positive results and seroprevalence were det ermined from ELISA assays.

| Location<br>(sampling period) | Species | Number of<br>distinct<br>individuals | Number of samples<br>(pre-covid/2021/2022) | Number positive<br>results<br>(pre-covid/2021/2022) | Seroprevalence %<br>(pre-covid/2021/2022) |
| --- | --- | --- | --- | --- | --- |
| Trois-Fontaines (2010-2022) | Roe deer | 539 | 942 (832/29/81) | 17 (16/1/0) | 1.8 (1.9/3.5/0.00) |
| Chizé (2010-2022) | Roe deer | 557 | 965 (817/29/119) | 23 (15/0/8) | 2.4 (1.8/0/6.7) |
| Aurignac (2019-2022) | Roe deer | 180 | 205 (97/49/59) | 7 (4/2/1) | 3.4 (4.1/2.0/1.7) |
| Gardouch (2019-2022) | Roe deer | 9 | 18 (6/4/8) | 0 (0/0/0) | 0.0 (0.0/0.0/0.0) |
| Domaine of Praillebard (2022) | Roe deer | 4 | 4 (0/0/4) | 0 (-/-/0) | 0.0 (-/-/0.0) |
| Vercors Vie Sauvage (2022) | Fallow deer | 16 | 16 (0/0/16) | 1 (-/-/1) | 6.3 (-/-/6.3) |
|  | Red deer | 9 | 9 (0/0/9) | 0 (-/-/0) | 0.0 (-/-/0.0) |
|  | Sika deer | 5 | 5 (0/0/5) | 1 (-/-/1) | 20.0 (-/-/20.0) |
| **Total** | **Roe deer** | **1,289** | **2,134 (1,752/111/271)** | **47 (35/3/9)** | **2.2 (2.0/2.7/3.3)**<br>95%CI: [1.6; 2.9]<br>([1.4; 2.8]/[0.6; 7.7]/[1.5; 6.2]) |
|  | **Fallow deer** | **16** | **16 (0/0/16)** | **1 (-/-/1)** | **6.3 (-/-/6.3)**<br>95%CI: [0.2; 30.2]<br>(-/-/[0.2; 30.2]) |
|  | **Red deer** | **9** | **9 (0/0/9)** | **0 (-/-/0)** | **0.0 (-/-/0.0)**<br>95%CI: [0.0; 33.6]<br>(-/-/[0.0; 33.6]) |
|  | **Sika deer** | **5** | **5 (0/0/5)** | **1 (-/-/1)** | **20.0 (-/-/20.0)**<br>95%CI: [0.5; 71.6]<br>(-/-/[0.5; 71.6]) |
|  | **All species** | **1,319** | **2,164 (1,752/111/301)** | **49 (35/3/11)** | **2.3 (2.0/2.7/3.7)**<br>95%CI: [1.7; 3.0]<br>([1.4; 2.8]/[0.6; 7.7]/[1.8; 6.4]) |

### 2.2 Sera collection

Sera were collected in these populations during 2,134 capture events, involving 1,289 different individuals. Each individual was sampled one to nine times during the study period (2010-2022), with 874 individuals sampled only once. All samples were collected each year in winter between December (year *t-1)* and March (year *t*), except at GA where sampling occurred until May. Sera were immediately stored at -80°C after collection on the field and during transportation to the laboratory, where they were stored at -20°C until analyses.

### 2.3 Nasal swabs collection

For direct virus detection, nasal swabs were collected in 2022 during the roe deer capture-mark-recapture programs from TF (n = 49), CH (n = 95) and AU (n = 61). Nasal swabs were immediately stored at -80°C after collection and during transportation to the laboratory where they were immediately tested upon arrival.

### 2.4 Additional sampling on other cervid species in other locations

Additionally, we sampled cervids in other locations: the Vercors Vie Sauvage (VVS) private reserve located in eastern France (44°57’N, 5°14’E; Figure 1, green star) and three departments of central France (Allier, Cher and Loire; Figure 1, blue areas).

We sampled 30 sera from other cervids captured in 2022 in the VVS, including 16 European fallow deer, five sika deer and nine red deer (Table 1). In the three central France departments, nasal swabs were also collected on hunted roe deer (Allier: n = 3, Cher: n = 29, and Loire: n = 4), and on red deer in one department (Cher: n = 3) (Figure 1, blue areas).

We defined two periods, either before COVID-19 emergence in France (pre-COVID-19 emergence: 2010-2020, n = 1,752, ‘pre-Cov’ hereafter) or after COVID-19 emergence (post-COVID-19 emergence: 2021-2022, n = 382, ‘post-Cov’ hereafter).

### 2.5 Initial ELISA serological screening

Serum samples were first screened using the ID Screen® SARS-CoV-2 Double Antigen Multi-species ELISA kit (ID Vet, France). This kit is designed for the detection of antibodies (IgG, IgM) against the nucleocapsid (N-protein) of SARS-CoV-2, and thus does not allow to discriminate between past and recent infection. The optical density (OD) was measured spectrophotometrically at 450 nm (Safas MP96, Monaco) following the manufacturer’s instructions. The seropositivity threshold was S/P% ≥ 60, with *S/P* = 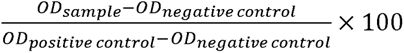

### 2.6 Confirmatory seroneutralization assay

To confirm or refute positive ELISA results, the corresponding samples were further analysed using seroneutralization as described in Pozzetto *et al.* [28]. Briefly, Murine leukemia virus-based pseudoparticles carrying a green-fluorescent protein (GFP) reporter pseudotyped with SARS-CoV-2 spike protein (SARS-CoV-2pp) were used to measure the neutralizing antibody activity in sera. A sample of *ca.* 10^3^ pseudoparticles was incubated with a 1/100 dilution of serum for 1 h at 37 °C before infection of the Vero-E6R cells (ATCC CRL-1586). The percentage of GFP-positive cells was compared to cells infected with SARS-CoV-2pp incubated without serum to set the percentage of neutralization. The positive control used was a rabbit monoclonal neutralizing antibody targeting the receptor-binding domain of the SARS-CoV-2 (2019-nCoV) spike protein (Sino Biological, 40592-R001), with a 100-fold dilution, as used in Pozzetto *et al.* [28] and Fritz *et al.* [29]. As negative control, we used a subset of 30 samples that were negative to ELISA tests, corresponding to the pre-pandemic period [29].

### 2.7 Direct detection

Nasal swabs (n = 244) were analysed using reverse transcriptase polymerase chain reaction (RT-PCR) technique. In an experimental study on white-tailed deer, RT-PCR on nasal swabs allowed to detect SARS-CoV-2 viral RNA at least 22 days post-infection, while infectious virions could only be detected for up to six days post-infection [30]. Briefly, automated extraction instruments (KingFisher Flex 96 instrument by Thermo Fisher Scientific, Waltham, MA, USA, or the Auto Pure 96 instrument by Allsheng Instruments CO, Hangzhou, China) were used to extract viral RNA from swabs using the BioExtract Premium Mag (Biosellal, Dardilly, France). Subsequently, SARS-CoV-2 RNA detection was performed by qRT-PCR with the Bio-T kit TriStar COVID-19 (Biosellal, France), which targets the envelope gene (E) of Sarbecoviruses (among which SARS-CoV-1, SARS-CoV-2), and RNA-dependent RNA polymerase (RdRp) in the Orflab gene, according to the manufacturer’s instructions. The AriaMx Real-Time PCR Thermal Cycler System (Agilent Technologies, Santa Clara, CA, USA) was used for amplification. The condition consisted of 1 cycle of 20 min at 50 °C, 1 min at 95 °C, followed by 40 cycles of 10 s at 95 °C, and 45 s at 60 °C. The results were analysed using the Agilent Aria 1.6 software, in which a cycle threshold value (Ct-value) < 40 for all target genes was defined as a positive result.

### 2.8 Analysis of seropositivity in roe deer

Only populations with positive samples were included in the following analyses: AU, CH and TF. Firstly, S/P% values distribution was inspected using Shapiro-Wilk test of normality and the number of components was assessed in a mixture of multinomials model based on AIC (Akaike Information Criterion) and BIC (Bayesian Information Criterion).

Then, we analysed the pattern of ELISA results to better understand the origin of seropositive signals. Under the hypothesis of SARS-CoV-2 emergence in roe deer populations following introduction from humans, a higher seroprevalence was expected in the post-COVID-19 human emergence period compared to the pre-COVID human emergence period. To test this hypothesis, Chi² tests (alpha = 0.05) were run to compare seroprevalence between the different periods for the three populations together and separately. The two periods were either: before COVID-19 emergence in France (pre-COVID-19 emergence: 2010-2020, n = 1,752, ‘pre-Cov’ hereafter) or after COVID-19 emergence (post-COVID-19 emergence: 2021-2022, n = 382, ‘post-Cov’ hereafter). We used samples of the first period as controls to test for potential cross-reaction with other coronaviruses circulating in these populations. We included the year 2020 in the pre-emergence period because sampling occurred between December 10th 2019 and March 12th 2020, before the virus spread across France and the lockdown was enforced on March 17th [31].

However, because animal manipulation was reduced in 2021 as a consequence of the implementation of social restrictions during the pandemic, the virus could have emerged in roe deer populations later than 2021 (i.e. in 2022). Thus, considering a possible delayed exposition of the roe deer population, we ran a second test comparing the 2010-2021 vs. 2022 periods. We thus defined two different periods: either before a delayed COVID-19 emergence in France (pre-delayed COVID-19 emergence: 2010-2021, n = 1,863, ‘pre-dCov’ hereafter) or after a delayed COVID-19 emergence with relaxed social restrictions (post-delayed COVID-19 emergence: 2022, n = 271, ‘post-dCov’ hereafter). Models with pre-Cov vs. post-Cov periods as well as pre-dCov vs. post-dCov periods were built.

All statistics were done with R version 4.5.0 [32] in Rstudio 2025.09.1 [33] and with the *readr* 2.1.5 [34] and *mixtools* 2.0.0.1 [35,36] packages.

## 3 Results

### 3.1 Serological results

A total of 47 out of 2,134 ELISA tests (2.2% [95%CI: 1.6; 2.9]) performed on roe deer sera had positive results for anti-SARS-CoV-2 antibodies against the N-protein (Table 1). It included samples collected during both the pre-Cov (35 positives: 2.0%) and post-Cov (3 positives: 2.7% and 9 positives: 3.3%, in 2021 and 2022, respectively) periods (Table 1). Among the 47 positive results, 17 out of 942 (1.8% [1.1; 2.9]) were found in TF (pre-Cov: 16 out of 832 (1.9% [1.1; 3.1]); 2021: 1 out 29 (3.5% [0.1; 17.8]); 2022: 0 out of 81 (0.0% [0.0; 4.5])), 23 out of 965 (2.4% [1.5; 3.6]) in CH (pre-Cov: 15 out of 817 (1.8% [1.0; 3.0]); 2021: 0 out of 29 (0.0% [0.0; 11.6]), 2022: 8 out of 119 (6.7% [3.2; 14.0])) and 7 out of 205 (3.4% [1.4; 6.9]) in AU (pre-Cov: 4 out of 97 (4.1% [1.1; 10.2]]); 2021: 2 out of 49 (2.0% [0.5; 14.0]); 2022: 2 out of 59 (1.7% [0.4; 11.7])). The 47 positive sera originated from 46 individuals from CH, TF, and AU (Table 1), including 28 individuals sampled and tested only once, seven tested twice, six tested thrice, four tested four times and one tested five times. During the follow up of the 19 positive individuals tested multiple times, both appearance and disappearance of antibodies were evidenced, as 13 seronegative individuals were found seropositive in the following years (six on the next year, seven two years or more later), while 11 negative results were observed on samples that were previously positive (four on the next year, seven two years or more later). Only one individual was tested positive twice during two different years (2016 and 2018), but no sample was available in-between (*i.e.* 2017). Twenty positive results were from individuals in their first year.

The S/P% values did not follow normal distribution (Shapiro-Wilk test: *p* < 10^-3^). It rather followed a bimodal distribution according to the mixture of multinomials model assessment (both AIC and BIC), as expected when a proportion of the population have been exposed to a pathogen and displays a high antibody concentration. Observed positive results ranged from 60.7 to 824.4%. Additionally, one roe deer was tested twice during the same year (with a three-week interval). Both samples from this individual were positive and observed S/P% declined from 328% to 120% between the two samples (only the first observation was kept in the analyses).

Although seroprevalence was higher in the post-COVID-19 emergence period (delayed or not) than before SARS-CoV-2 emergence, this difference, when considering all roe deer populations together, was not statistically significant (pre-Cov vs. post-Cov Chi² test: χ² = 1.71, df = 1, *p* = 0.19; pre-dCov vs. post-dCov Chi² test: χ² = 1.51, df = 1, *p* = 0.22). However, this difference was significant at CH with or without considering an emergence delay (pre-Cov vs. post-Cov Chi² test: χ² = 5.41, df = 1, *p* = 0.02, odds ratio = 3.05; pre-dCov vs. post-dCov Chi² test: χ² = 8.96, df = 1, *p* < 0.01, odds ratio = 3.99).

Among the sera from other cervids inhabiting the VVS reserve, ELISA tests performed on one fallow deer out of 16 (6.3% [0.2; 30.2]) and one sika deer out of five (20.0% [0.5; 71.6]) yielded seropositive results (Table 1).

SARS-CoV-2 seroneutralization assays were performed on nine positive and 15 negative roe deer samples of the post-COVID-19 emergence period, and on four positive and 15 negative roe deer samples of the pre-COVID-19 emergence period. All seroneutralization assays, including the 13 carried out on ELISA-positive samples, yielded negative results.

### 3.2 Direct virus detection

RT-PCR carried out on nasal swabs collected during winter 2022 on 205 captured roe deer (of which 10 were positive with the ELISA assay) and 36 hunted roe deer and three hunted red deer, did not reveal any positive result.

## 4 Discussion

This study is the first long-term individual monitoring of serological status of wild-living cervids for SARS-CoV-2. Our survey intended to detect, at the individual or population levels, an increase in seropositivity or seroprevalence following the emergence of SARS-CoV-2 in France. In contrast with most studies investigating SARS-CoV-2 seroprevalence in European deer species, the present one encompasses longitudinal data (*i.e.* repeated measurements of some individuals), captive vs. wild individuals and individuals exposed to direct handling by humans. It also includes 2022 data, whereas most studies were limited to 2020 and 2021 [16–18]. In fact, only the most recent investigations detected cases of SARS-CoV-2 in deer populations [20–22], highlighting the need to continue SARS-CoV-2 surveillance of European deer because new variants are continuously emerging [37,38].

Interpreting serological results in terms of viral infection is not straightforward. SARS-CoV-2 ELISA tests indicated an overall seroprevalence of 2.20% in roe deer. We also observed positive results in other cervids from VVS despite a low sample size (one out of 16 fallow deer and one out of five sika deer). However, all sera submitted to the seroneutralization assay, and all swabs tested for direct detection of the virus by RT-PCR in 2022 gave negative results. Furthermore, seroprevalence was significantly different before and after the SARS-CoV-2 emergence in humans, whether considering a delayed transmission to roe deer or not, only in one out of three population (*i.e.* CH). This indicates that sarbecoviruses, among which SARS-CoV-2, are unlikely to circulate in the studied populations of roe deer.

The absence of confirmation of SARS-CoV-2 antibodies by seroneutralization assay is consistent with other studies conducted in Germany, Austria and the UK on roe deer, red deer, fallow deer, sika deer, Reeves’s muntjac, and Chinese water deer [16,18,19,21]. Negative seroneutralization assays despite positive ELISA tests have also been reported in wild boars (*Sus scrofa*), red foxes (*Vulpes vulpes*) and jackals (*Canis aureus moreoticus*) [19,39]. It can be explained by an inadequate specificity of the ELISA test allowing cross-reaction with antibodies directed to other closely-related viruses [40]. Also, while ELISA used in this study detect antibodies targeting SARS-CoV-2 nucleoprotein, the seroneutralization assay detects neutralizing antibodies targeting spike glycoprotein. Previous studies showed a poor correlation between antibodies targeting nucleoproteins and neutralizing antibodies in SARS-CoV-2-infected patients [41]. This could also explain why a serum can score positive in a nucleoprotein-based ELISA assay, but negative in a seroneutralization assay. Such results underline the importance of not relying on ELISA tests alone before drawing any conclusions.

Nonetheless, these positive results (with S/P% values up to 824.4% and a bimodal distribution) suggest that the ELISA test used might cross-react with one or several other coronaviruses circulating in the studied roe deer populations. Viruses close to bovine coronaviruses (BCoVs) have already been found in red deer and sika deer [42], and are potential candidates that could have been detected by our ELISA analyses. Positive serological results were observed in all populations of roe deer, except DP and GA for which only four and 18 sera were tested, respectively. If a virus of bovine origin is involved, one would expect a lower seroprevalence in the two isolated roe deer populations of CH and TF compared to the free-roaming roe deer population of AU exposed to cattle. Our results do not show such pattern and in principle would not support the BCoVs hypothesis.

The circulation of phylogenetically close coronaviruses transmitted by humans (*e.g.* HCoV-OC43 [43]) is also possible. In case of a human transmitted coronavirus, at least a similar seroprevalence in captive roe deer in GA (experimental manipulation all year long) and roe deer in TF or CH (captures during winter) would be expected, but no positive individual was observed in GA (acknowledging, however, a small sample size). Thus, our results do not support the human coronavirus hypothesis either.

Other possibilities would include: (*i*) long-lasting BCoVs infection maintenance in the isolated populations since the enclosure of the areas, (*ii*) repeated introduction of BCoVs by infected animals entering the enclosure by some gaps in the fences, which is possible for roe deer or another susceptible species like wild boar, or (*iii*) exposure of cervid populations to various coronaviruses from other wildlife species present in the studied areas. Indeed, we cannot exclude the eventuality that coronaviruses are transmitted to roe deer by bats, rodents, rabbits or hedgehogs (*e.g.* alpha- and beta-coronaviruses [3,44]), or that another, still unknown, coronavirus is circulating in cervid populations, as suggested elsewhere [19,43]. The seropositivity of a fallow deer and a sika deer, which were not in such close contact with human or cattle than studied roe deer, supports the idea of an exposure of wild cervid populations to at least one cross-reacting coronavirus.

However, as the virus(es) responsible of the observed seroprevalence is unknown, one cannot exclude the possibility that the different populations are exposed to distinct viruses (*e.g.* BCoVs in AU, other coronaviruses in CH and TF). More investigations are required before drawing any firm conclusion, for instance by using more specific tests to confirm or infirm the presence of BCoVs antibodies in ELISA positive roe deer sera [45], or conduct High Throughput Sequencing analyses on cervid samples to identify candidate viruses [46].

Among 11 positive individuals that were tested the following years, only one was found seropositive twice. Other individuals showing seropositive status were tested negative during the following years, including four individuals tested only one year later. These results are consistent with a short persistence of the antibodies (less than one year), but for at least several weeks (>3). Although a rapid decline of humoral response has been described, antibodies after natural SARS-CoV-2 infection in humans can persist for more than a year [47,48]. In white-tailed deer, it should last at least 13 months [49], but persistence in other species is not well known. Thus, according to available data in white-tailed deer, as persistence time is expected to be longer and juveniles are not expected to have a higher seroprevalence, our observations are not consistent with a SARS-CoV-2 emergence in the studied roe deer populations [49,50]. Still, we detected a significant difference between the pre- and post-COVID-19 emergence periods (delayed or not) in CH only, showing an increase in seropositivity following COVID-19 emergence. Data from coming years may clarify these temporal trends in the different populations. Nonetheless, without formally identifying the virus(es) causing the observed response, the current patterns remain difficult to explain. The different dynamics between populations might result from distinct circulating viruses or variants with specific dynamics [20,51] and that would need to be identified.

Transmission of the SARS-CoV-2 virus from human to roe deer is thus unlikely to have happened to date in the studied populations, contrary to what has been observed among white-tailed deer populations in North America [12–14]. Our results confirm the results of previous studies evidencing an absence of contact between the most common European cervids and the SARS-CoV-2 virus [16–19], except in densely populated areas [20–22]. The three European studies reporting deer infection cases comprise individuals living in suburban areas and parks highly frequented by humans and in direct contact with them [20–22], confirming the pattern observed in white-tailed deer in North America. In contrast, other European studies examined deer populations without close contact with humans [16–19]. More specifically, roe deer is usually not living in peri-urban areas thus contacts of human with roe deer are expected to be less common than with white-tailed deer. Moreover, hunting management differs between North America and Europe, resulting in drastically different spatial and temporal distribution of deer populations. White-tailed deer thrive in urban and suburban areas seasonally reaching high densities, while European populations are maintained locally and in rural areas [18]. In the present study, individuals were totally preserved from human contact, except during scientific handling during winter. However, due to the COVID-19 pandemic, all manipulations were conducted in compliance with public health measures, including the use of hand sanitizer and surgical masks. Therefore, it is likely that differences in deer infections with SARS-CoV-2 between North-America and Europe (and among European studies) result from a combination of factors such as close contact with humans, species ecology and high population densities, rather than from species-specific susceptibility only.

Also, the ACE2 protein, which interacts with the SARS-CoV-2 spike protein, is identical at all import sites between roe deer and white-tailed deer [18], however, roe deer appear to be less susceptible to SARS-CoV-2 infection [18,21]. These results together suggest a necessary complementarity of molecular and ecological factors for interspecific viral transmissions. Despite the low probability of roe deer, or other European cervids, to serve as a reservoir host for SARS-CoV-2, the circulation of one or several other coronaviruses, maybe BCoVs, in roe deer populations might raise the question of a possible virus cross-recombination leading to a new virus variant. Further studies should be carried out to identify coronaviruses circulating in wild European cervids, and to continue testing roe deer populations to enable track temporal trends. Maintaining serological surveillance and implementing efficient virological surveillance to detect viruses in wildlife is necessary to evaluate the possibility of transmission between wild and domestic animals and humans [52].

## Conflict of interest disclosure

The authors declare that they have no conflict of interests relating to the content of this article.

## Data and code availability

All data and codes used for the present analyses have been deposited on the CIRAD dataverse and are publicly available: https://doi.org/10.18167/DVN1/AR6UDN [53].

## Fundings

This project was funded by the *Université de Lyon* (Project RESPOND and Doctoral grant), *VetAgro Sup* and the *Office Français de la Biodiversité* (OFB, project CNV-REC-2019-08). Funders were not involved in the study design, the collection, analysis and interpretation of data, the writing of the report and the decision to submit the article for publication.

## Ethics and authorisations

The research presented in this article was done according to all institutional and/or national guidelines. The protocol of capture and blood sampling of roe deer is under the authority of the *Office Français de la Biodiversité* (OFB) and was approved by the Director of Food, Agriculture and Forest (Prefectoral order 2009-14 from Paris). All procedures were approved by the Ethical Committee of Lyon 1 University (project DR2014-09, June 5th, 2014). The Trois-Fontaines population is part of the long-term Studies in Ecology and Evolution (SEE-Life) program of the CNRS. The data from Aurignac population came from a long-term study approved by ethical committee n°115 and by French Ministry of Higher Education and Research (Apafis#39320).

## Authors contributions

**Study conceptualisation:** GB and EGF

**Methodological design:** GP, GB and EGF

**Field sampling:** GP, LL, HV, VB, NC, YC, PR, MP, JFL, CP, BR, FD, RG, GB and EGF

**Immunological assays:** GP, LL and BR

**Seroneutralization assays:** VL

**RT-PCR screening:** AK

**Study coordination and supervision:** GB and EGF

**Further supervision:** MP and JFL

**Funding:** JFL, GB and EGF

**Statistical analyses:** GP and LL

**Writing – original draft:** GP and LL

**Writing – first editing:** GB and EGF

**Writing – review and editing:** GP, LL, VL, AK, HV, VB, JFL, CP, BR, GB and EGF

## Acknowledgements

This work was conducted as part of a Ph.D. and postdoctoral projects funded by the *Université de Lyon*, the *Office Français de la Biodiversité* (OFB) and *VetAgro Sup*. We are grateful to all technicians, researchers and volunteers participating in the collect of data on all sites (Trois-Fontaines, Chizé, Aurignac, Gardouch, Domaine de Praillebard) and laboratory analyses. The Trois-Fontaines population is part of the long-term Studies in Ecology and Evolution (SEE-Life) program of the CNRS. We also thank the *Asociation Protection des Animaux Sauvages* for the Vercors Vie Sauvage samples and hunters from Allier, Cher and Loire for samples from hunted wild cervids.

## Notes

https://doi.org/10.18167/DVN1/AR6UDN

### Competing Interest Statement

The authors have declared no competing interest.

### Summary of Updates

Version updated following peer review and acceptance in Royal Society Open Science. A previous version of this preprint was recommended by PCI Infections. Changes include: - putting the current study within the context of research in European cervids regarding SARS-CoV-2 investigations - The removal of the modelling part of the manuscript - A comparison of the North-American and European situations

